# Synergic microRNAs suppress human glioblastoma progression by modulating clinically relevant targets

**DOI:** 10.1101/2025.03.27.645741

**Authors:** Silvia Rancati, Rui C. Pereira, Michele Schlich, Stefania Sgroi, Silvia Beatini, Letizia La Rosa, Lidia Giantomasi, Roberta Pelizzoli, Clarissa Braccia, Carlotta Spattini, Kiril Tuntevski, Amanda Lo Van, Meritxell Pons-Espinal, Annalisa Palange, Adriana Bajetto, Antonio Daga, Andrea Armirotti, Tullio Florio, Paolo Decuzzi, Davide De Pietri Tonelli

## Abstract

Glioblastoma (GBM) is a highly aggressive brain tumor characterized by therapy-resistant glioma stem-like cells (GSCs) and extensive infiltration into surrounding brain tissue. MicroRNAs (miRNAs) are post-transcriptional regulators of oncogenic pathways, but their tumor-suppressive function is frequently lost in GBM. This study explores a multimodal therapeutic approach by restoring a combination of miRNAs to exploit their synergistic effects against GBM. Using patient-derived GBM cells cultured under stem cell-permissive conditions, we demonstrate that miRNA restoration reduces tumor growth, limits invasiveness, stemness and enhances sensitivity to temozolomide. *In vivo* studies in an orthotopic xenograft mouse model of GBM confirm the therapeutic efficacy and low toxicity of the nanoformulated miRNAs, following local injection. Multi-omics and computational analyses on different GBM subtypes reveal that these miRNAs synergistically suppress tumor-promoting extracellular matrix interactions, particularly through the collagen pathway, and downregulate genes associated with GBM progression. The identified miRNA targets correlate with glioma grade and poor patient prognosis, further underscoring their therapeutic potential. These findings highlight the promise of combinatorial miRNA therapy as a novel strategy for GBM treatment and suggest new molecular targets for theragnostic development.

## INTRODUCTION

Glioblastoma (GBM) is the most common and aggressive malignant brain tumor in humans, with a median survival of approximately 14.5 months, a figure that unfortunately has not improved over the past two decades^1^. Recurrence is the unavoidable outcome of GBM, likely driven by therapy-resistant, heterogeneous glioma stem-like cells (GSC) subpopulations, which are reminiscent of neural stem cells (NSC) ^2,3^. This is in line with the hypothesis that GBM may originate from the malignant transformation of NSCs ^3–5^. GSCs are predominantly quiescent and infiltrate the non-neoplastic brain parenchyma, complicating surgical resection, and resulting in the development of therapy-resistant clones following standard-of-care treatments ^6,7^. Moreover, as confirmed by the limited success of therapeutic strategies aimed at one or few targets, GBM is unlikely to be cured by targeting one or few factors, in line with the notion that cancer is not a unique disease caused by a single mutation or epigenetic alteration ^8^. Thereby the identification of effective multimodal therapies, ideally aimed at GCSs depletion and preventing infiltration, is an urgent need for this malignancy.

MicroRNAs (miRNAs) are small noncoding RNAs that by post-transcriptional repression of the majority of protein coding genes in mammals ^9^ maintain proper transcriptional homeostasis ^10,11^. Since the original report of miRNA dysregulation in cancer ^12^, several studies have shown that this is also the case in glioma ^13,14^, in line with the aberrant activation of genetic programs in this malignancy ^15^. Unsurprisingly, miRNAs are regarded as promising tools and targets for cancer therapy ^16^. However, despite various miRNA-based drugs have been included in clinical trials, many of those targeting single miRNAs have failed, indicating a need for further development ^17^. Conversely, the use of miRNA combinations has recently gained momentum as a therapeutic strategy in neuro-oncology ^13,18–22^, in line with the functional redundancy (*i.e.,* synergism) of miRNAs in physiological NSCs ^23^.

We previously identified a pool of eleven pro-neurogenic miRNAs (henceforth “the pool”) that sustains adult neurogenesis through the synergistic modulation of different targets in NSCs ^24^. Since the rescued miRNA pool was sufficient to restore impaired neurogenesis in miRNA-depleted NSCs, which resemble GSCs, we hypothesized that these miRNAs might be effective against GBM. Here, we study the expression of the eleven miRNAs across various glioma grades and GBM subtypes in patients and demonstrate the efficacy of the pool as a gene-therapy against GBM. Finally, by multi-omics analysis we infer the mechanisms underlying the tumor-suppressive functions of the pool in different GBM subtypes and uncover that these miRNAs suppress clinically relevant targets in glioma.

## RESULTS

### 1. Underexpression of the eleven miRNAs in gliomas of different grades

To correlate the abundance of the eleven miRNAs (*i.e.* miR-124-3p; miR-127-3p; miR-134-5p; miR-135a-5p; miR-139-5p; miR-218-5p; miR-370-3p; miR-376b-3p; miR-382-5p; miR-411-5p; miR-708-5p) with human glioma grade, we analyzed public datasets of Low-Grade Glioma (LGG) from The Cancer Genome Atlas (TCGA), available in DIANA-miTED database ^25^ and of non-neoplastic human brain, as control ^26^. Ten of the eleven miRNAs (except miR-134-5p) are significantly underexpressed in grade II and grade III gliomas compared to controls (Figure 1A). Because grade IV glioma (i.e. GBM) was not available in this dataset, abundance of the eleven miRNAs was quantified by small RNA sequencing (RNA seq) and quantitative PCR (qPCR) in cultures obtained from surgical samples of patients with glioma grade III (GG3) and GBM (GBM1-2)^27^. This analysis indicated that the eleven miRNAs are underexpressed in two grade IV gliomas (GBM1, classified as proneural and GBM2 classified as mesenchymal subtype, respectively; *data available upon request, manuscript in preparation*) and in one GG3 ^27^, compared to non-neoplastic controls (Figure 1B, statistics in Supplemental Table 1). QPCR confirmed that eleven and nine of the miRNAs were underexpressed in GBM1 and GBM2, respectively, compared with control brain (Figure 1C). These results, indicating variations in the abundances of the eleven miRNAs across distinct GBM grades and subtypes, are in line with the heterogeneity of this malignancy and suggest that underexpression of these miRNAs is likely associated with gliomagenesis and/or tumor progression.

**Fig. 1.**
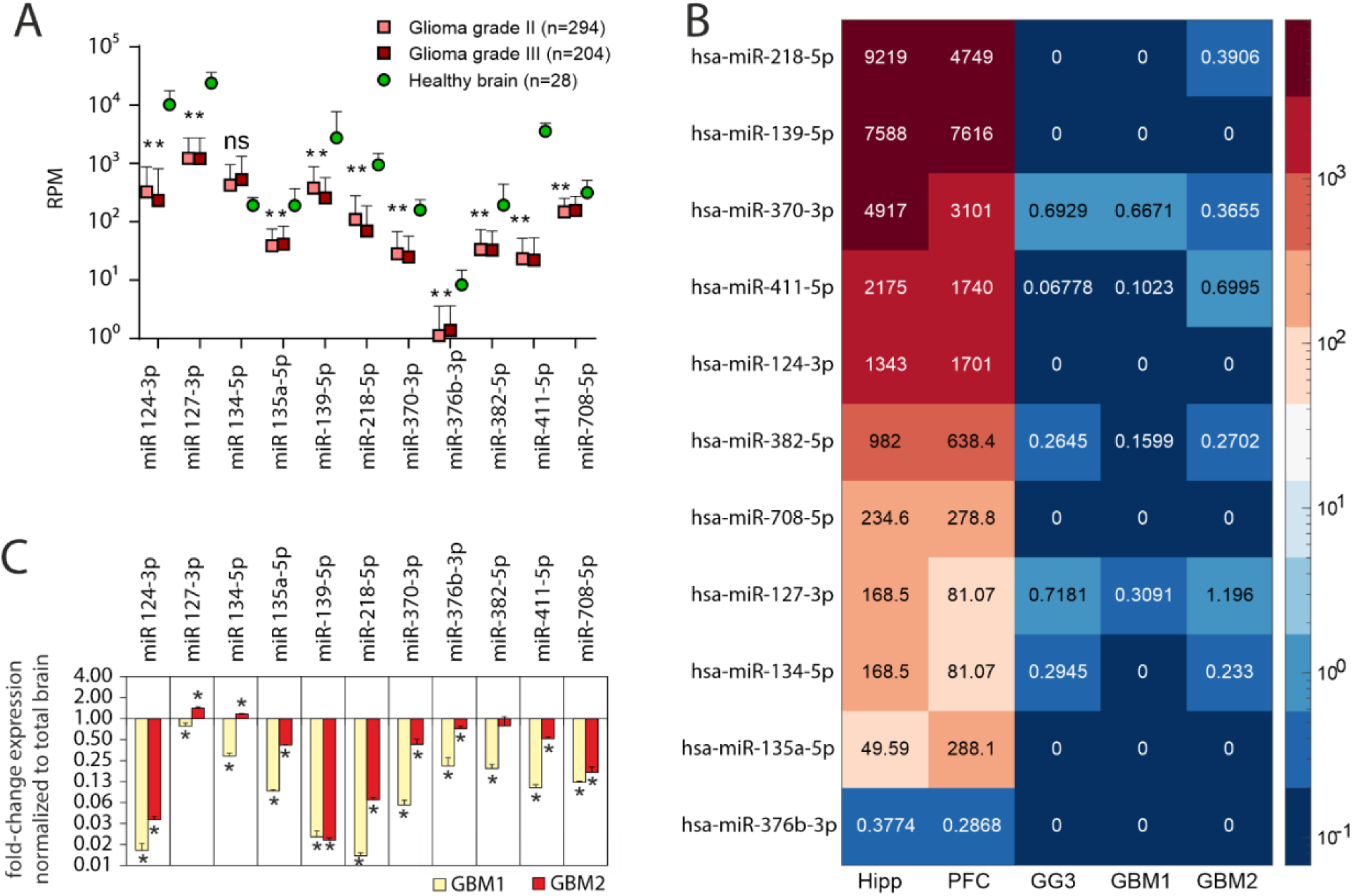
Underexpression of the eleven miRNAs in gliomas of different grades. **A)** Levels of the 11 miRNAs in glioma samples (from TCGA-LGG cohort: n=294 grade II and n=204 grade III glioma) and non-neoplastic brain tissue (n=28). Data expressed as Reads Per Million (RPM); (average ± SD; *p<0.001, unpaired t-test) ^25^. **B)** Heatmap of small RNA seq showing levels of the 11 miRNAs in the cell cultures from two GBM patients (GBM1, 2), in Glioma Grade III cells (GG3), and non-neoplastic human hippocampus (Hipp) and prefrontal cortex (PFC) from GEO database accession# PRJNA752352 and GSE181520). Data are the mean TPM in n=3 biological triplicates. **C)** Quantification (qPCR) of abundances of the 11 miRNAs in the GBM1, 2 cultures normalized to n=1 sample of non-neoplastic brain RNA (data are the mean ΔΔct ± St.Dev of n=1 experiment in technical triplicate; two-tailed t-test *p<0.05).

### 2. The pool significantly reduces GBM growth, invasiveness, and increases sensitivity to temozolomide (TMZ)

To test whether the pool has an effect against GBM, we transfected synthetic mimics of the eleven miRNAs in equimolar concentrations (*i.e.,* ∼25 nM each of the eleven miRNAs), or a scrambled RNA mimic (Control) to a final concentration of 250 nM in both GBM cultures and maintained them as monolayers. Restoration of the eleven miRNAs was confirmed by qPCR five hours (h) after transfection (*see black bars in* Figure 5A). To investigate the phenotypic effect of the pool, we performed a scratch-wound assay and used label-free automated live imaging to quantify invasion (*i.e.,* cells invading a layer of Matrigel), migration (i.e., cells migrating by adhesion on plastic) (Figure 2A) and growth (*i.e.,* cell confluence, Figure 2B) of both GBM cultures. The pool significantly reduced the growth of both GBMs (Figure 2B), and invasiveness in GBM2, but not GBM1 (Figure 2A). In contrast, the pool only slightly delayed migration of monolayer cultures of both GBM (Figure 2A). Hence, the pool can significantly reduce GBM growth and invasiveness *in vitro*, suggesting it may be more effective in mesenchymal rather than in the proneural subtype.

**Fig. 2.**
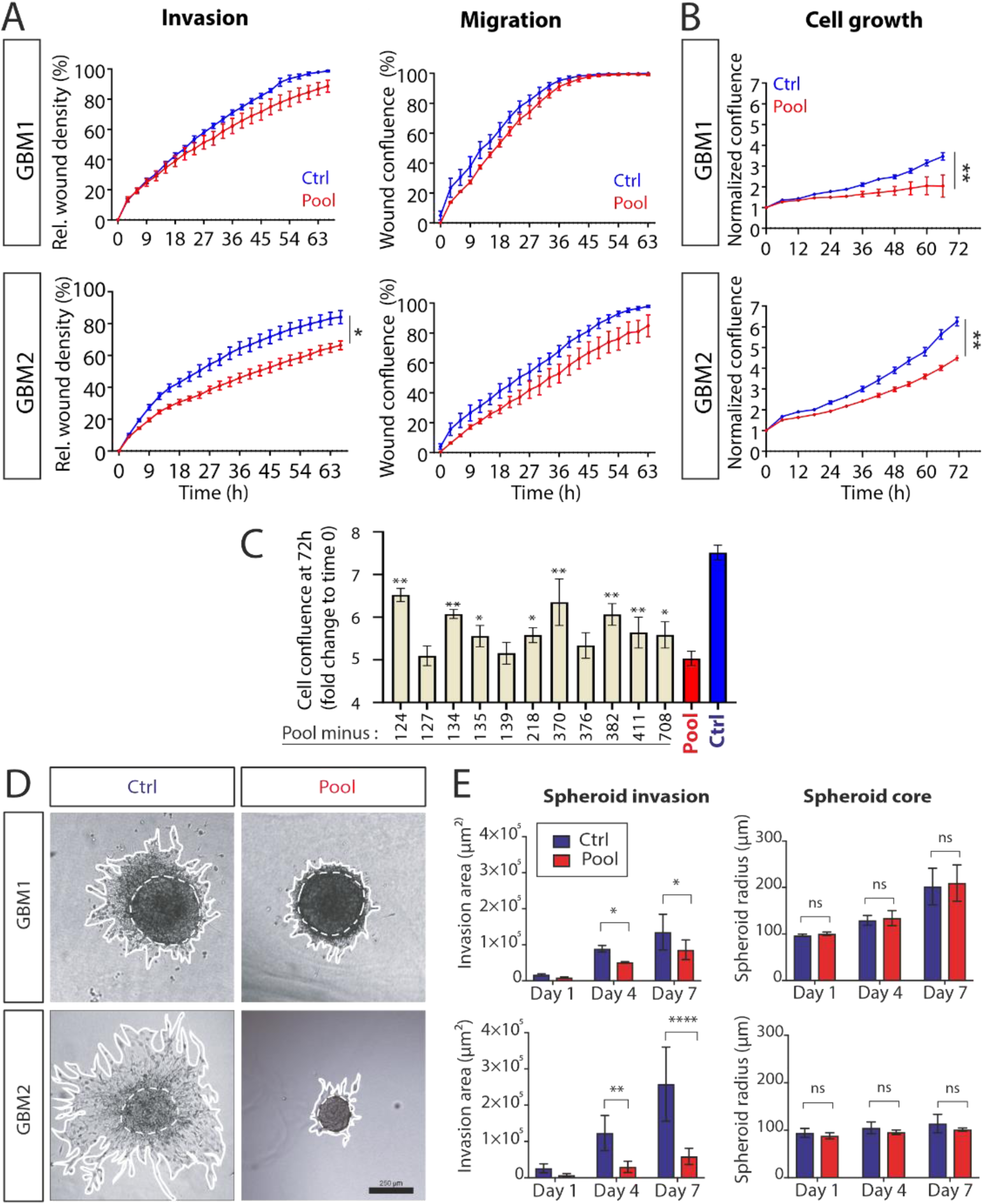
The pool significantly reduces GBM growth and invasiveness. **A)** Scratch wound assay quantification of invasion or migration of scrambled RNA (ctrl) and pool-transfected GBM 1, 2 (data are mean ± St.Err. of n=1 experiment, in triplicate, two-way ANOVA interaction p value (time x treatment): *<0.05). **B)** Quantification of growth of scrambled RNA (“ctrl”) and pool-transfected GBM1, 2 (mean ± St.Err. of n=1 experiment; five replicates; two-way ANOVA interaction p value (time x treatment) **p<0.005). **C)** Growth of GBM2 cells transfected with scrambled RNA (“ctrl”), pool or sub-pools (i.e., each missing one miRNA). Plotted data is mean ± St.Dev of cell confluence at 72 h normalized on time 0 (i.e. 24 h after transfection). The missing miRNA is indicated in the x-axis; (n=1 experiment, five replicates; asterisks mark statistical significance of each condition versus pool: *p<0.05, **p<0.005; one-way ANOVA and multiple comparisons; all the tested conditions are statistically significant when compared to scrambled). **D)** Representative images showing spheroids from scrambled RNA- (ctrl) and pool-transfected GBM1 and GBM2 after 7 days in 3D collagen. **E)** Quantification of spheroid cores (*dashed circles*) and invasion area *(delimited by the dashed and the white line*). Scale bar 250 μm (Data are expressed as means ± St.Dev. of n > 3 independent experiments, each with n > 5 spheroids per condition; Two-way ANOVA and multiple comparisons: *p<0.05; **p<0.005 ****p < 0.0001).

Given the observed heterogeneity in the miRNA abundances across different GBM subtypes (Figure 1), we asked whether all the miRNAs are required to elicit the above effects in GBM. To address this question, we quantified growth of GBM2 cells upon the transfection of the pool with all the eleven miRNAs (Pool) or eleven combinations, each missing one of the miRNAs (Pool minus “miR-*x”*), versus scrambled RNA (Figure 2C, Control). Quantification of cell confluence at 72h indicated an attenuation of the effect of the pool on GBM2 growth when one miRNA was not restored (Figure 2C). This result is in line with our previous findings in NSC ^24^, confirming that this pool acts synergistically also in GBM.

To validate the effects of the pool in a GBM model which better mimics the 3D features and interaction of the tumor with the microenvironment ^28^, we formed spheroids of GBM from cells that were pre-transfected with the pool or scrambled RNA, and then followed their growth and infiltration upon embedding them in collagen hydrogels. In contrast to cells cultured in monolayers, the pool did not alter growth of the spheroid core area (dashed circles in Figure 2D) compared to scrambled control, whereas the invasion area (*i.e*. the surface occupied by cells protruding out of the core, *white lines in Figure 2D*) was significantly reduced in both GBM cultures upon transfection with the pool; again, a stronger inhibition of infiltration was observed upon pool treatment in the mesenchymal GBM2, compared to proneural GBM1 subtype (Figure 2E).

To assess the efficacy of the pool in combination with chemotherapy, pool- or scrambled-transfected GBM1 and GBM2 were cultured in presence of different TMZ concentrations (1:2 dilutions ranging from 1500 to 46 µM), and growth documented by imaging for up to 72h (Supplemental Figure S1). Remarkably, the pool increased TMZ sensitivity of GBM1 (but not of GBM2, *not shown*) compared to scrambled RNA (Supplemental Figure S1).

Together, these results demonstrate that the pool has a tumor-suppressive role in GBM, primarily by reducing invasiveness, and suggest that these miRNAs may likely enhance GBM sensitivity to TMZ.

### 3. The pool represses the collagen pathway and markers of GSCs subpopulation

To gain a mechanistic insight, we sought to identify genes and pathways regulated by the pool in GBM. Therefore, we performed Liquid Chromatography/Mass Spectrometry (LC/MS) proteomics and RNAseq, to shed light on the effects of the eleven miRNAs on the proteome and transcriptome, respectively. Pool- and scrambled-transfected spheroids of both GBM subtypes, a condition previously shown to preserve GSCs ^29^ and tumor-initiating properties ^27^, were harvested after 3 or 7 days for RNAseq and proteomics, respectively. Untargeted proteomics identified ∼ 4,000 unique proteins expressed in each of the two pool- transfected or scrambled-transfected GBM cultures (3962 in GBM1; 4206 in GBM2). The pool significantly altered 8.8% and 11% of proteins in GBM1 and GBM2 (FDR<0.05), respectively, compared to scrambled-transfected controls (Supplemental Table 2). Principal component analysis (PCA) of the proteomics data indicated a clear segregation of the two GBM cultures along the PC1 (Supplemental Figure S2), in agreement with the heterogeneity that characterizes this malignancy ^6^ and with the different gene expression profiles of the two samples ^30^. Consistently with the observed tumor-suppressive effects of the pool, we found an evident segregation of the proteomes of the pool- and scrambled-transfected GBM samples along PC2 (Supplemental Figure S2). Analogously, the pool significantly altered 4% and 5.9% transcripts (FDR<0.05) in GBM1 and 2, respectively (Supplemental Table 3). These results indicate that the pool modulates a substantial portion of the proteome and transcriptome in different GBMs.

Given that miRNAs act mostly as repressors^16^, to identify the targets underlying the tumor suppressive functions of the pool, we focused on genes and proteins downregulated in both GBM cultures (Figure 3A). As expected for such heterogeneous malignancy, most of the downregulated genes were different between the two cultures, and only 37 genes and 37 proteins were commonly downregulated among them (Figure 3A, *top*; *the green box shows “common targets”, i.e. downregulated in both GBM at transcript and protein level*). Notably, despite the inherent differences between the two GBM subtypes, Gene Ontology (GO) analysis indicated that the pool suppresses similar pathways in both GBM models, particularly cell-adhesion, cell membrane mediators of the extracellular matrix (ECM) interaction and production, with several pathways being shared between proteomics and transcriptomics datasets (Figure 3A, *bottom*). Gene Set Enrichment Analysis (GSEA) of transcriptomics data (Supplemental figure S3), highlights that “Epithelial Mesenchymal transition” (EMT) is among the top downregulated pathways by the pool in both GBMs, together with “TNF-alpha signaling via NFKβ” and “TGFβ signaling”, two main inducers of EMT, and involved in GBM invasiveness and therapy-resistance ^31^. PLOD3, MMP14, NQO1 and several members of collagen IV and integrin family were suppressed by the pool in both GBM subtypes at both the transcript and protein levels (Figure 3A, *Top, “common targets”*). Notably, Protein-protein interaction (PPI) analysis indicated a high degree of functional redundancy between the proteins downregulated by the pool (Figure 3B), as indicated by their inclusion in a network with significantly enriched number of interactions. K-means analysis performed by STRING algorithm clustered the proteins into 3 main groups, whose functions can be summarized as: ECM adhesion and remodelling (*cluster A*), collagen production and metabolism (*cluster B*), Aldehyde Dehydrogenases (*cluster C*) (Figure 3B). Interestingly, the latter family of enzymes is overexpressed in GBM and has been associated with therapy-resistance, invasiveness and stemness ^32^. Among the downregulated transcripts/proteins/pathways, we noted an enrichment in ECM-interacting components, and particularly collagen-related pathways: indeed, 13 of the 37 proteins (*red names* in Figure 3B) are involved in collagen synthesis, post-translational modification, remodeling, recycling or binding, suggesting that the pool might inhibit tumor ability to synthesize and/or interact with this ECM. In agreement, the pool inhibited GBM2 cell adhesion (Figure 3C) on collagen, and invasiveness of both GBM subtypes in this ECM (*see Figure 2D,E*). This evidence indicates that the tumor-suppressive action of the pool is primarily mediated by the modulation of cell-to-cell and cell-to-ECM adhesion, particularly by suppressing genes of the collagen pathway in both GBM subtypes.

**Fig. 3.**
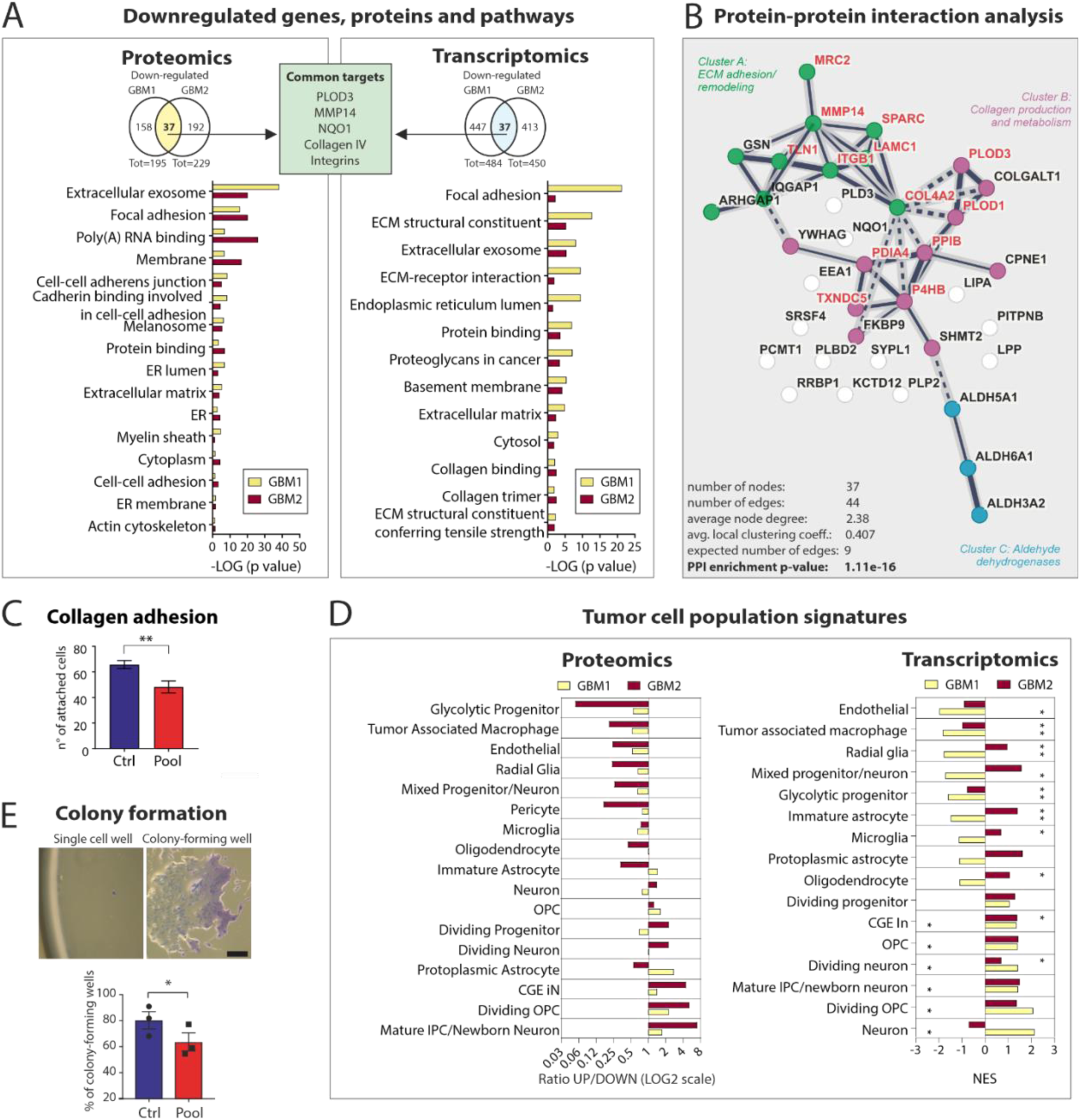
Multi-omics analysis, assessment of adhesion and colony formation in GBM cultures after restoration of the pool. **A)** Proteomics and transcriptomic analysis of scrambled RNA and pool-transfected GBM1,2. *Top*, Venn diagram showing the number of proteins or genes downregulated by the pool in each GBM (FDR <0.05) and the ones in common (LC-MS proteomics: n=5 samples, 7 days after transfection; RNAseq transcriptomics: n=3 samples, 3 days after transfection). The green square highlights the common putative miRNA targets significantly downregulated by the pool compared to control in both GBM1, 2 at the protein/transcript levels. *Bottom*, pathway analysis (Gene Ontology) of the 37 proteins downregulated by the pool in both GBM cultures (For each GO term is plotted the negative Log of Benjamini-adjusted p-value. **B)** STRING protein-protein interaction (PPI) analysis of the 37 downregulated proteins in common between GBM1, 2. **C)** Quantification of scrambled or pool-transfected GBM2 cells attached on collagen IV-coated coverslips after 5 minutes (n=1 experiment, in 5 replicates; Two-tailed unpaired t-test, **p<0,005). **D)** Analysis of the relative enrichment of tumor cell population signatures (clusters as defined in ^2^) in proteomics and transcriptomics of GBM1, 2 upon pool transfection. Proteomics: Ratio of the number of up- to down- regulated proteins from each cluster (ratio <1: downregulated by the pool). Transcriptomics: Normalized Enrichment Score (NES) of each cluster obtained from GSEA analysis of pool- versus control-transfected GBM (NES <0: downregulated by the pool; *FDR q-value<0.005). **E)** Quantification of colony-positive wells 2 weeks after single-cell seeding of pool- or scrambled-transfected GBM2 cells; data is normalized to the number of colony-positive wells of untransfected cells (n=3 independent experiments, at least 30 wells/condition; two-tailed, paired t-test; 12 *p<0,05).

To address whether the miRNA pool modulates GSC subpopulations in the two GBMs, we classified the transcripts/proteins that were differentially expressed upon pool transfection using clusters of GBM cell subpopulations from previous single-cell RNAseq studies ^2,33^. Interestingly, markers of GSC subpopulations (*glycolytic progenitor* and *radial glia*) ^3^ and those associated to tumor-promoting features (*tumor-associated macrophages*) were less represented in both pool-transfected GBM subtypes (Figure 3D). The pool also downregulated markers related to vasculogenic potential of GSCs (*endothelial, pericyte)* (Figure 3D), which can trans-differentiate into vascular endothelial cells ^34^. To ascertain whether the reduction in the abundance of stemness markers reflected a depletion of GSC subpopulation, we seeded individual pool- or scramble-transfected cells and performed a clonogenic assay on Matrigel-coated coverslips by quantifying the generation of new colonies. Two weeks after pool transfection, we found a significantly lower number of colony-forming wells, compared to scrambled control (Figure 3E). This result indicates that restoration of the pool leads to a reduction of the GSC subpopulation.

Collectively, this evidence indicates that the pool reduces invasiveness and stemness in GBM cultures, and strongly suggests that the modulation of cell-to-cell and cell-to-ECM adhesion, particularly the downregulation of the collagen pathway, represents a common mechanism underlying the tumor-suppressive functions of the pool in different GBM subtypes.

### 4. Implementation of a synthetic nanocarrier for *in vivo* intratumor delivery of small RNAs in GBM

To implement a delivery strategy suitable for testing the pool efficacy *in vivo*, we engineered a nanocarrier by encapsulating equimolar concentrations of the eleven miRNA mimics in Lipid Nanoparticles (LNPs) via microfluidic-based mixing (Figure 4A). Notably, a similar formulation has been approved by the US Food and Drug Administration (FDA) to deliver siRNA to hepatocytes in the treatment of transthyretin amyloidosis ^35^.

**Fig. 4.**
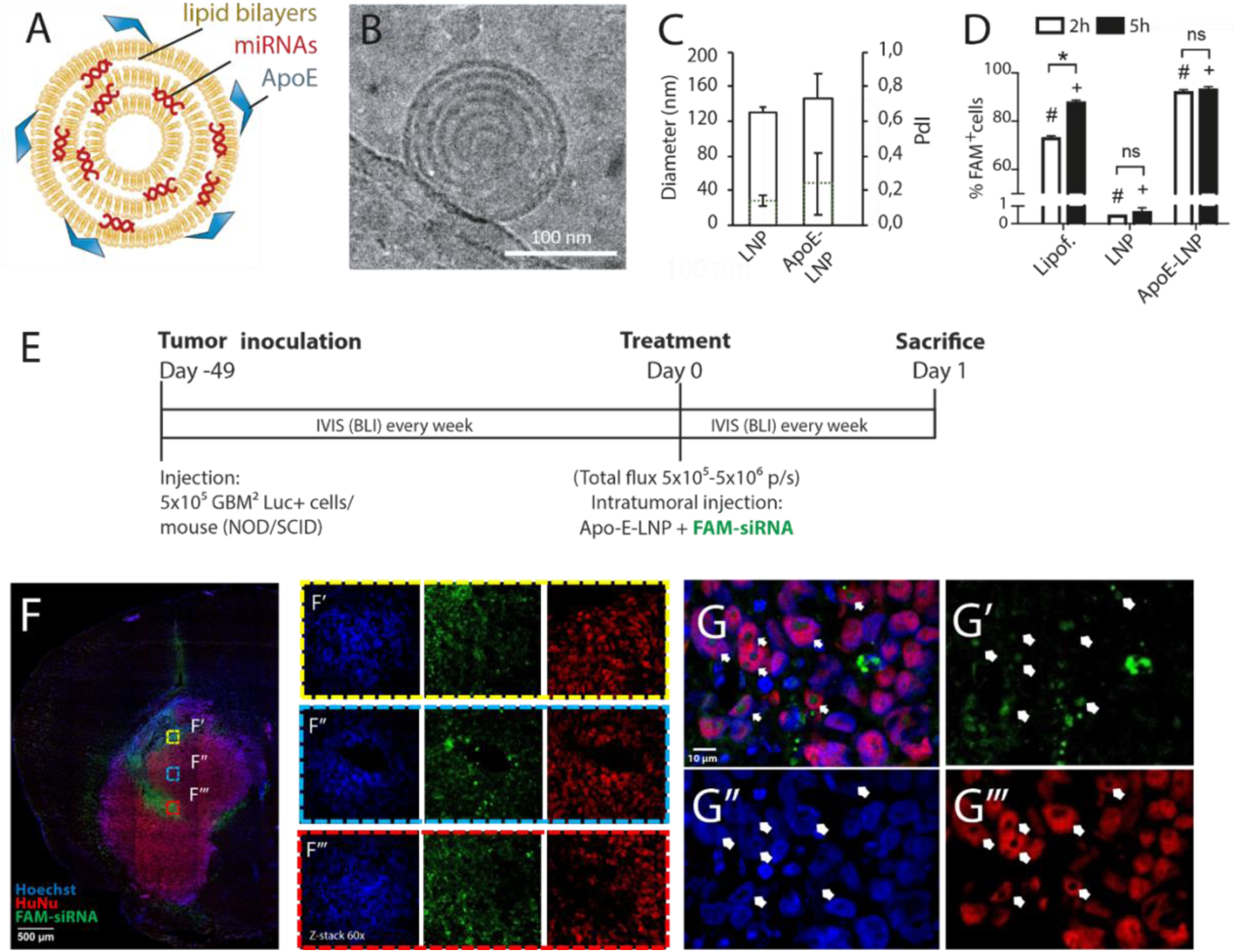
*In vitro* characterization of the nanocarrier and *in vivo* biodistribution of the small RNA cargos, upon intratumor injection in preclinical mouse model with orthotopic GBM xenotransplants. **A)** Scheme of the ApoE-LNP nanocarrier. **B)** Cryo-electron microscopy showing one representative image of LNP (scalebar 100 nm). **C)** Diameter and polydispersity index (PdI) measurement of LNP with or without ApoE coating (mean ± St.Dev of at least 4 independent batches measurements). **D**) Percentage of U87MG cells positive for the FAM-labelled siRNA fluorescence 2h or 5h upon Lipofectamine2000 (positive control), LNP, or ApoE-LNP incubation, as determined by Flow cytometry (n=2 replicates) (# and + marks conditions statistically different among them; two way ANOVA and multiple comparisons). **E)** Experimental scheme of GBM2 orthotopic xenotranplantation in NOD/SCID mice and intratumour injection of ApoE-LNP loaded with FAM-siRNA. **F)** Representative images of FAM-siRNA (green) distribution in the tumor mass (Hoechst, *blue*; FAM-siRNA, *green*; Human Nuclei – HuNu–, *red*; Objective 10x, Scale bar 500µm) 24 h after intratumor injection of FAM-siRNA-loaded ApoE-LNP. **F’-F’’’)** Higher-magnification showing the distribution of the three signals inside the tumor mass (indicated by the boxed regions in F; z-stack 60x). Note the green signal intensity is similar between the three tumor areas at different distances from the injection site/tumor mass center. **G)** Confocal images showing merged (G) and separate channels (G’-G’’’) of FAM-siRNA (green) inside the human GBM cells (HuNu, *red;* Hoechst, *blue*). Note that GBM cells appear bigger than the (HuNu-negative) mouse cells. Objective 60x, scale bar 10 µm.

The charge of the eleven miRNAs was neutralized by ionizable lipids, mixed with neutral lipids and PEGylated lipids to generate particles of ∼120 nm diameter (Figure 4B). To overcome the low uptake of uncoated LNPs by cancer cells ^36^, Apolipoprotein E (ApoE), which is the ligand of the lipoprotein receptor overexpressed in several cancer cell types including GBM ^37^, was adsorbed on the surface of the nanoparticles (Figure 4A). Cryo-electron microscopy characterization of the LNPs showed a multi-lamellar structure (Figure 4B), in line with previous reports for similar particles. The LNP and ApoE-LNP hydrodynamic diameters were 127 nm and 135 nm, respectively (*bars* in Figure 4C). A low polydispersity index (PdI) was observed (*dotted lines* in Figure 4C), indicating a homogenous size distribution of the particles for both preparations. To ascertain the coating with ApoE, we measured the zeta potential of the LNPs. As expected, the zeta potential of ApoE-LNPs was negative compared to uncoated LNPs and thus consistent with the negative charge of this protein (Supplemental Figure S4).

To test whether the nanocarriers could deliver miRNAs into GBM cells *in vitro*, we loaded FAM-labelled siRNA (same size of miRNA mimics) in LNPs with- or without ApoE-coating and administered them to a human GBM cell line (U87MG). As a positive control, we transfected the same siRNA with Lipofectamine. As expected, FAM-labelled siRNA loaded in LNPs was not uptaken efficiently by the cells as indicated by flow cytometry analysis at both 2 and 5 hours after cell treatment (Figure 4D). Conversely, FAM-labelled siRNAs were successfully uptaken by the cells when loaded into ApoE-coated LNPs, even more efficiently than with Lipofectamine, at both time points after treatment (Figure 4D). Confocal imaging of GBM cell line treated with ApoE-LNPs confirmed the intracellular localization of FAM-labelled siRNA (Supplemental Figure S4).

To test the toxicity *in vivo*, we injected the nanoformulated FAM-labeled siRNA in the brain of healthy mice and no tissue damage, or cell death, was observed in the area surrounding the injection (*data not shown*), in agreement with the known low neurotoxicity of ApoE-LNPs ^38^. To test the biodistribution and the intracellular delivery of miRNAs via ApoE-LNPs, we implemented a xenotransplant mouse model by orthotopic injection of human GBM cells (*i.e.,* GBM2) constitutively expressing the luminescent reporter luciferase (Luc^+^) in the striatum of NOD-SCID mice. Tumor formation and growth were monitored for the subsequent 49 days by the IVIS Spectrum *in vivo* imaging system (IVIS). FAM-labelled siRNAs loaded in ApoE-LNPs were administered locally by a single intratumor injection, and brains were harvested 24 hours after for the analyses (Figure 4E). Fluorescence confocal microscopy imaging of immunostained sections across the brain of NOD-SCID mice showed FAM-fluorescence (Figure 4F, *green*) uniformly distributed in several regions of the engrafted tumor, identified by the positivity for the human-specific nuclear antigen HuNu, (Figure 4F, *red*). Higher magnification of the engrafted tumors confirmed the intracellular delivery of the FAM-labeled siRNAs in GBM cells (Figure 4G). Hence, a single injection of the nanoformulation allows efficient intratumor distribution and intracellular delivery of a synthetic siRNA in GBM *in vivo*.

### 5. A single intratumor administration of the nanoformulated pool reduces GBM progression *in vivo*

To test whether the ApoE-LNPs could simultaneously deliver the eleven miRNAs to GBM cells, we loaded either the eleven miRNAs or the scrambled RNA in the ApoE-LNPs. The encapsulation efficiency was 79.5 ± 5.3 % for the miRNA pool and 86.3 ± 2.0 % for the scrambled. QPCR of GBM2 cultures treated with pool-loaded ApoE-LNPs confirmed the restoration of all the eleven miRNAs, compared to cells treated with scrambled RNA-loaded ApoE-LNPs, with the same efficiency of Lipofectamine (Figure 5A).

To assess the phenotypic effect of the nanoformulated pool in the preclinical model of GBM xenotransplantation *in vivo*, 42-49 days after implantation of Luc^+^ GBM2 cells in the striatum of NOD/SCID mice, we injected intratumorally pool- or scramble- loaded ApoE-LNPs. Growth of GBM tumor xenografts was monitored for two weeks after the injection of the nanoformulated RNAs by IVIS- mediated quantification of luciferase signal, and brain harvested for further analyses (Figure 5B-D). *In vivo* stability of the nanoformulated pool was assessed by quantifying levels of the hsa-miR-376b (the only human-specific miRNA of the pool) by qPCR with human-specific primers (Supplemental Figure S5). Hsa-miR-376b levels were significantly increased in the GBM tumor 24 h after local injection of the ApoE- LNPs loaded with the pool, compared to GBM injected with control ApoE-LNPs (Figure 5C). However, at 14 days post injection, levels of the hsa-miR-376b returned to almost basal (Figure 5C). Assuming similar kinetics for all the miRNAs of the pool, this result suggests that the nanoformulated miRNAs are stable for less than two weeks after intratumor injection *in vivo*. Remarkably, a single injection of the ApoE-LNPs pool significantly inhibited the rate of GBM growth for two weeks following the treatment, compared to the group of mice injected with scrambled-loaded ApoE-LNPs (Figure 5E), in line with the stability of the miRNA mimics *in vivo*. This result confirms the tumor-suppressive efficacy of the pool in a preclinical model of human GBM.

**Fig. 5.**
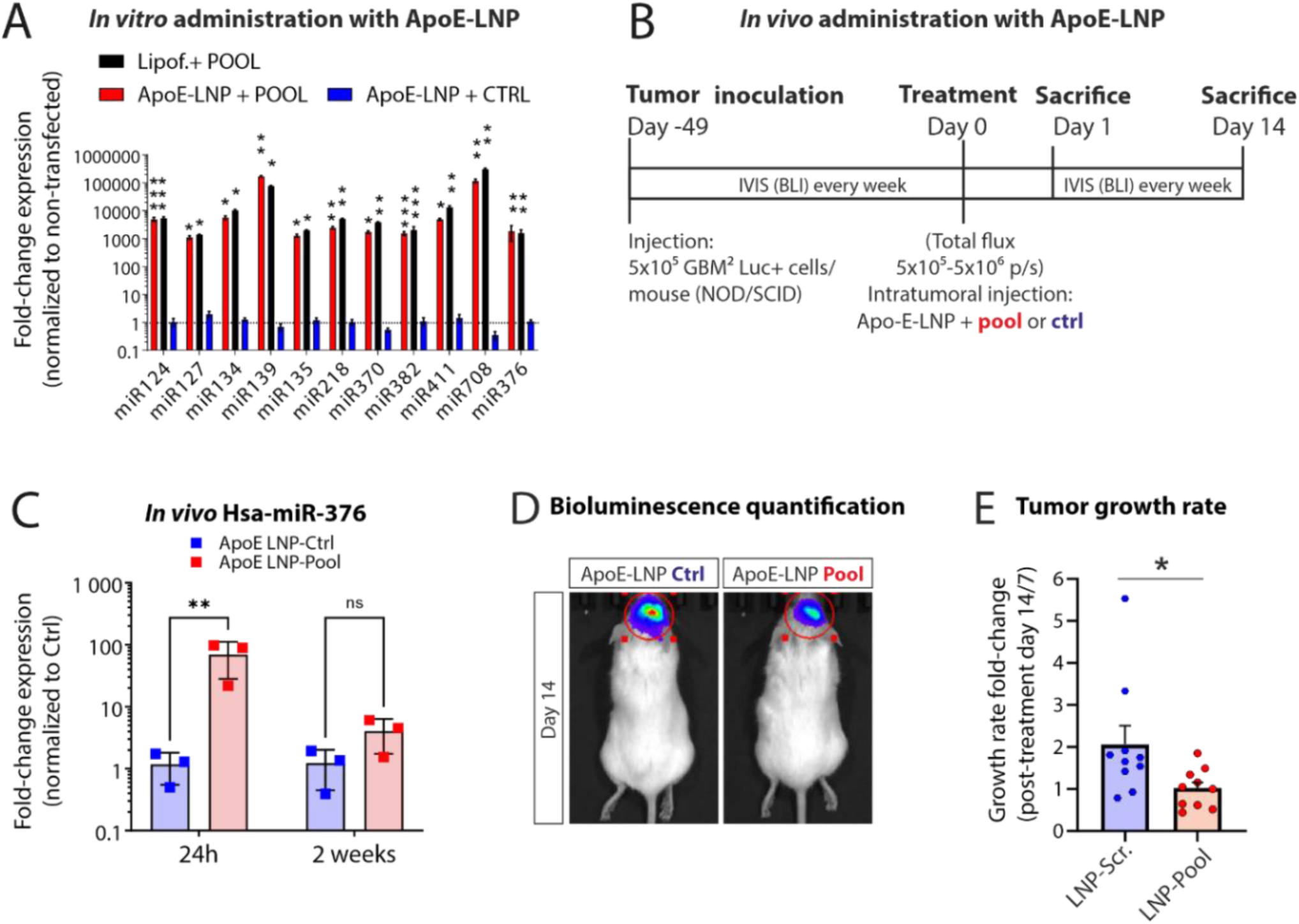
*In vivo* stability and therapeutic efficacy of the miRNAs upon a single intratumor injection of nanoformulated pool. **A)** QPCR quantification of the eleven miRNAs levels 5h after *in vitro* delivery of the pool with Lipofectamine, or pool- or scrambled control (ctrl)-loaded LNPs, coated or uncoated with ApoE. (Data are expressed as fold-change expression and normalized to untreated condition; ΔΔct mean ± St.Dev; ApoE-LNP samples are n=2 biological replicates, qPCR samples are analyzed in technical triplicate). **B)** Experimental scheme of the *in vivo* xenotransplantation and intratumor injection of the nanoformulated pool. **C)** QPCR quantification of the levels of hsa-miR376b in xenotransplanted GBM after intratumor injection of nanoformulated pool or scrambled RNA. Data are expressed as fold-change expression normalized to ApoE-LNP control at each time point (ΔΔct mean ± St.Dev; n=3 animals per condition; **p<0.005, two-way ANOVA and multiple comparisons). **D)** Representative images of xenotransplanted mice 14 days after intratumor injection of nanoformulated pool or scrambled RNA as assessed by IVIS bioluminescence. **E)** Tumor growth rate represented as ratio of the total flux (p/s) assessed by IVIS bioluminescence at day 14 / day 7 upon intratumor injection of the nanoformulated pool or scrambled RNA (n=10 animals; Data are the mean ± SEM; *p=0.03, unpaired two-tailed t-test).

### 6. Clinically relevant targets underlie the tumor-suppressive functions of the pool *in vivo*

To validate the mechanisms underlying the tumor-suppressive effect of the pool in xenotransplanted GBM, two weeks after a single intratumor injection of pool- or scramble-loaded ApoE-LNP, we performed bulk transcriptomics of tumor tissue (Figure 6A). To identify potential miRNA targets, we focused on transcripts that were downregulated following *in vivo* injection of the pool and compared them with those identified *in vitro* (see Figure 3). This analysis confirmed eight genes that were downregulated in both *in vivo* and *in vitro* conditions (Figure 6B). Notably, five out of the eight downregulated genes encode for proteins involved in cell adhesion, motility or ECM interaction in GBM, or other cancer types (*i.e.,* PCDH18 ^39^, COL3A1 ^40^, ITGA5 ^41^, FN1 ^42^ and ROBO1 ^43^), and the other three encode for either transcriptional or epigenetic regulators involved in EMT (*i.e.,* FOXS1 ^44^), stemness (*i.e.,* BCL9L ^45^), or GBM growth (*i.e.,* JMJD6 ^46^). The GO pathway analysis of the downregulated genes upon pool injection *in vivo* (Figure 6C) showed again cell- and ECM-adhesion pathways, largely overlapping with the ones from *in vitro* transcriptomics (*see Figure 3 and Supplemental S3*). Concomitantly, this analysis revealed several neural, synaptic and neuro-developmental GO terms downregulated upon pool administration *in vivo* (Figure 6C). Indeed, neural-specific pathways and synaptic genes have been associated to GBM infiltration by several reports, in line with the fact that GSCs hijack the neuronal genetic programs to invade in the brain parenchyma ^47,48^. These results confirm that repression of the tumor-ECM interaction and EMT-related pathways underlies the tumor suppressive functions of the miRNA pool in GBM *in vivo*.

**Fig. 6.**
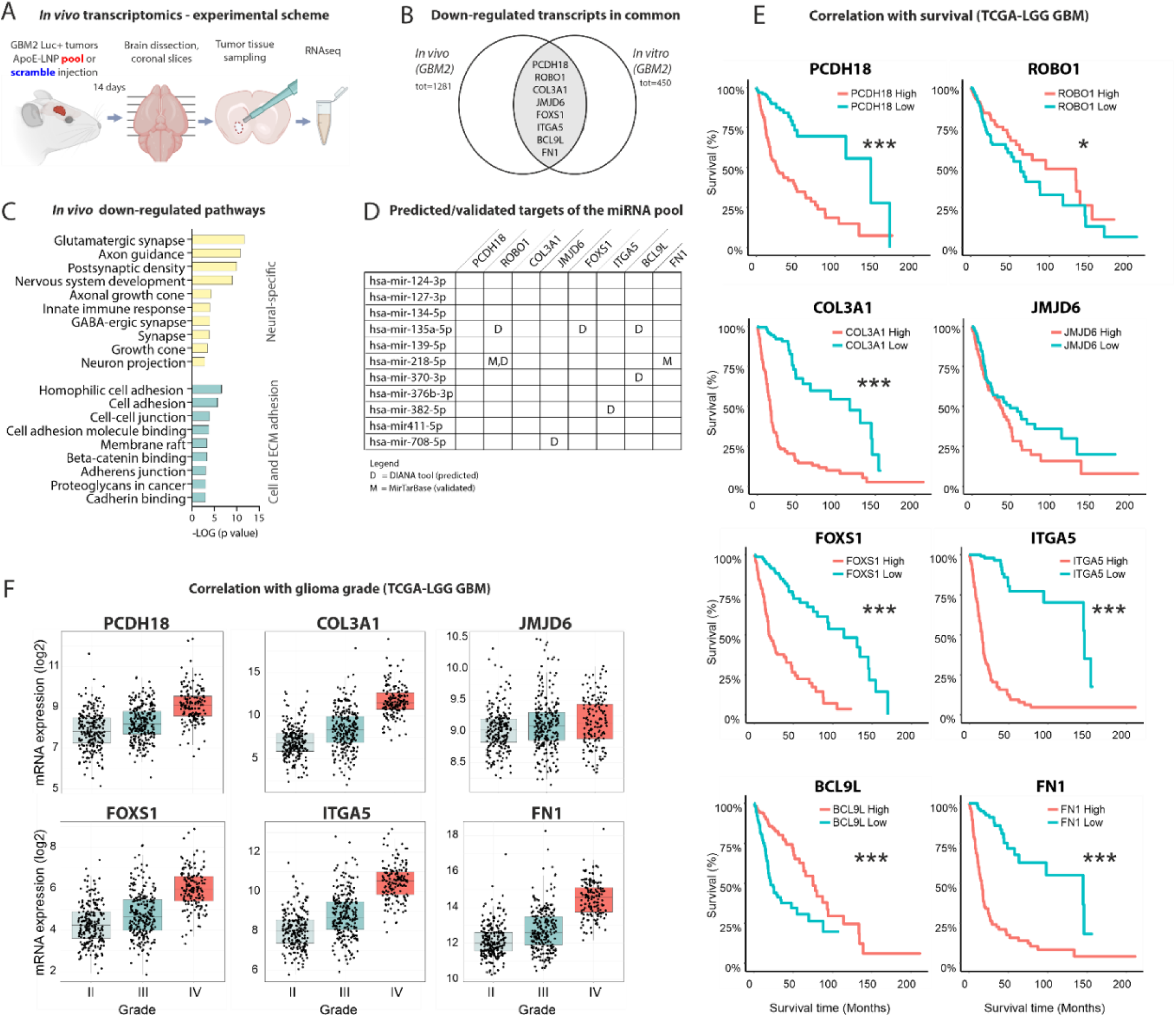
Genes, pathways and targets suppressed by the pool *in vivo* are prognostically relevant in GBM patients. **A)** Experimental scheme of the *in vivo* transcriptomics experiment (n=5 animals per group). **B)** Venn diagram showing the transcripts downregulated in GBM upon *in vitro* and *in vivo* administration of the pool from RNAseq data. **C)** Pathway analysis (GO enrichment) of transcripts downregulated *in vivo* upon administration of the miRNA pool. The most statistically significant GO terms and KEGG pathways are grouped in two macro-areas: nervous system cell development and morphology (“neural-like”, *yellow bars*) and cell-/ECM- adhesion (*blue bars*). Only terms with FDR <0.05 were selected, negative Log of p-value is plotted. **D)** Table summarizing the miRNA target analysis of the *in vivo/vitro* common downregulated genes, showing predicted (DIANA-microT, combining miRBase 22.1 and MirGeneDB 2.1 databases) and validated targets (miRTarBase). **E)** Kaplan-Meier survival curves of TCGA-LGG GBM patients for the 8 downregulated genes (cutoff: high vs low, log-rank p value: ***<0.001, *<0.05). **F)** Correlation between expression of the 8 downregulated genes and glioma grade from TCGA-LGG GBM dataset. (n=620, statistics in Suppl. Table S3).

To determine which of the downregulated genes were directly repressed by the miRNAs, we examined miRNA–mRNA interaction databases. We found that the majority (six out of eight) of the repressed genes (*see Venn diagram* in Figure 6C) are predicted or validated targets of one or more miRNAs from the pool (Figure 6D), suggesting they are likely regulated by these miRNAs. Finally, to understand the clinical relevance of the eight genes suppressed by the pool in GBM, we retrieved clinical information from TCGA-LGG GBM database. Interestingly, the expression of seven out of eight genes positively correlates with reduced survival in patients (Figure 6E), in line with previous reports in GBM ^39–42^. Moreover, six of the downregulated genes positively correlate with glioma grade (Figure 6F), being significantly overexpressed in GBM compared to grade II and III gliomas. These results confirm that the pool represses key genes involved in mechanisms that are aberrantly upregulated in GBM, highlighting new targets suitable for future theragnostic development.

## DISCUSSION

This study highlights the synergistic potential of pro-neural miRNAs as a multimodal therapeutic strategy for GBM, one of the most lethal cancers in humans. We restored the expression of eleven miRNAs that are underexpressed in gliomas and likely involved in GBM initiation and progression. Our results show that this miRNA cocktail significantly reduces GBM growth, invasion, and stemness in both *in vitro* and *in vivo* models. A single intratumor injection of the nanoformulated miRNA pool significantly slowed human GBM progression in a preclinical model. Hence, this study successfully addresses relevant problems underlying the poor prognosis of GBM patients.

Based on the existing literature, this study is the first to demonstrate the efficacy of a “fully synthetic” formulation of different miRNAs to treat human GBM *in vivo.* Unsurprisingly, the effects were transient, in line with the synthetic nature of the formulation. Hence, we anticipate that multiple rounds of local administration will likely be required to improve the survival rate of the xenotransplanted animals. Intratumor injection of synthetic formulations of small RNAs is feasible in the clinical setting and may offer several advantages, including a potentially faster approval process compared to viral and/or cell-based vectors ^15,49^. Future studies should explore alternative delivery methods, such as biodegradable implants ^50^, or chemical modifications of the mimics to enhance stability of the miRNAs ^51^ to prolong therapeutic effects.

As related to the translational value of the miRNA synergism as a multimodal gene therapy, recent publications have convincingly demonstrated that restoring combinations of different miRNAs significantly improves anticancer efficacy compared to a single miRNA^13, 17–21^. However, so far polycistronic transcripts bearing different precursors were typically used to express combinations of miRNAs in GBM ^19,20^. This strategy still poses some challenges that complicate their clinical translation. Indeed, it does not guarantee a constant transcription of the polycistronic precursor, nor the equimolar expression of the different miRNAs in the tumor. Moreover, cleavage of hairpin-miRNAs from polycistronic precursors and miRNA maturation rely on endogenous biogenesis proteins such as Drosha and Dicer, often dysregulated in cancer^52^. Furthermore, when the polycistronic precursor is delivered by viruses, the tropism of this carrier may not allow equal targeting of the different cell subpopulations in GBM. Additionally, viral carriers may trigger immune responses, complicating their repeated administrations. While some of the limitations related to the viral transduction have been recently overcome by using exosomes as carriers ^20^, this approach relied on a plasmid-encoded polycistronic precursor to express multiple miRNAs, which may still suffer from the limitations of such constructs discussed above. Nonetheless, the path to the therapeutic use of multiple miRNAs for GBM is now wide open, and we are confident that one of the promising RNA delivery strategies currently in trials will soon be pivotal in translating the miRNA synergism approach in neuro-oncology^17^.

At the functional level, the multi-omics analysis indicates that the pool represses the collagen pathway and suggests that this may contribute to reducing the GSC niche, providing a mechanistic framework for the tumor-suppressive effects of these miRNAs. This is in line with the reduced adhesion and invasiveness of GBM cells upon restoration of the pool. Indeed, collagen is a crucial component of the basement membrane, the brain-specific ECM that surrounds blood vessels, a microenvironment that provides means to both GSC niche maintenance and GBM cell migration. In fact, GBM cells invade through the brain parenchyma along the vessels, and ability of the malignant cells to interact with collagen is a crucial determinant of invasiveness and maintenance of the stem cell niche ^53^. The precise cascade of events underlying the efficacy of the pool against the GSC subpopulation warrants further studies. Nonetheless, our results align with the notion that modulation of genes involved in ECM organization, cell-ECM adhesion, and promotion of cell migration occurs in GSCs as they transition toward “differentiated” glioma cells ^54^.

Finally, despite the inherent heterogeneity of GBM, we identified key genes and pathways to inhibit tumor progression. Because these genes positively correlate with poor prognosis and higher glioma grade in patients, we are confident they have the potential to serve as targets for future therapies and diagnostic development.

## MATERIALS AND METHODS

### Cell cultures and in vitro assays

High grade primary GBM cells (tumor ID GBM1: GSC#Ge003 1740R-48-11; Tumor ID GBM2: GSC#Ge023 1740R-48-17 ^30^; GG3: GBM5 from ^27^) were established and cultured as previously described ^27^. Unless otherwise specified, Matrigel-coated wells were covered with Matrigel diluted 1:200 in DMEM, which was incubated for 30’ at 37°C and removed before cell plating. U87MG (from ATCC) were grown as monolayer in EMEM supplemented with 2mM L-Glutamine and 10% FBS (Gibco). <u>Transfection:</u> Lipofectamine2000 (LifeTechnologies) was used to transfect the miRNA pool (25 nM of each miRNA mimic) or Scrambled siRNA as control (miRIDIAN, Dharmacon); in each subpools of 10 miRNAs the missing miRNA was compensated by 25 nM of scrambled RNA, to maintain equal final concentrations. <u>Migration/invasion assay:</u> 24 hours (h) after transfection, 30000 cells/well were seeded in Matrigel-coated ImageLock® (Sartorius) 96-wells plates, after additional 24 hours the scratch was produced with a wound maker (Sartorius). Cells were kept in complete medium (migration assay) or coated with a layer of diluted Matrigel (Corning, 1:40 in DMEM, Gibco) and let in the incubator for 20’ before adding complete medium (invasion assay). <u>Cell growth and TMZ:</u> 24 h after transfection, 5000 cells/well were plated in Matrigel-coated 96-well plates and imaged for 72 h with Incucyte® (Sartorius) using Phase Confluence mask analysis to obtain confluence normalized on time zero. TMZ powder (Merck) was dissolved in DMSO (100 mM stock) and diluted in complete medium to a concentration of 1500 µM. From this, serial dilutions (1:2) were prepared in complete medium. <u>3D cultures in Collagen hydrogels:</u> 24 h after transfection, 500 cells/well were seeded in an ultra-low attachment 96-wells plates (InSphero) for 24h. Spheroids were transferred into collagen type I hydrogels and cultured for 7 days. Hydrogel preparation: collagen type I solution (3 mg/ml, SigmaAldrich) was mixed with a PBS, EMEM (5x) with phenol red, 0.1 M NaOH and sterile water to yield a final collagen concentration of 2 mg/ml at pH of 7.4. <u>Cell adhesion on</u> <u>collagen:</u> 5000 cells/coverslip of control- or pool-transfected GBM cells were seeded 72 h after transfection on glass coverslips pre-coated with human collagen IV (Corning; 10 mg/mL), and let adhere for 5 minutes; then, they were washed and cultured for two hours in fresh medium, fixed with 4% PFA, stained with Hoechst (1:1000) and counted. <u>Clonogenic Assay:</u> GBM cells were transfected as indicated above. 24 h after transfection, single cells were seeded in Matrigel-coated 96-wells and colonies were imaged under light transmitted microscope for 2 weeks (w) with a 10X objective in an Olympus CKX41 inverted microscope. The colony-forming wells were counted and normalized on colony-forming wells of non-transfected condition. <u>Flow cytometry:</u> U87MG Cells were seeded at 8×10^4^ cells/well on 24-well plates. 24 h later, cells were treated with FAM-loaded-LNPs (100nM final FAM-RNA concentration) in complete medium. Transfection of the FAM-RNA (MISSION® siRNA, Merck) with Lipofectamine2000 was used as positive control. 2 and 5 h after LNP treatment, cells were analyzed with a FACs aria^TM^ IIIu cytofluorometer in PBS.

### Animal procedures

All care of animals and experimental procedures were conducted in accordance with IIT animal use committee and were approved by the Italian Ministry of Health. NOD/SCID female mice (Charles River, Italy) were group housed in ventilated cages in a climate-controlled animal facility (22 ± 2°C) and maintained on a 12 h light/dark cycle with *ad libitum* access to food and water. <u>Surgery</u>: Anesthetized animals (2% isoflurane/0.8% oxygen) were operated with a stereotaxic apparatus keeping them on a warming pad at 37°C. 3 µl of suspension containing 5×10^5^ human GBM cells (GBM2-Luc+) were injected into the striatum (coordinates, AP:1; ML: −2; DV: −3 from Bregma) by a syringe (Hamilton) connected to a pump; skin was sutured with absorbable wire, and mice were monitored until recovery. <u>Intratumor injection of nanoformulated miRNAs:</u> 42-49 days after xenotransplantation (*i.e.,* when the rate of tumor growth started to rise, as quantified by IVIS), 3 µL of ApoE-LNP loaded with FAM-labeled siRNAs, or pool or scramble RNA (corresponding to 1.35 nmol of RNA) were injected in each animal. The site of injection and procedure were the same used for GBM cells implantation. *See “LNP” methods in Supplemental Information*.

### Histology

24 hours after injection of FAM-labeled siRNAs loaded in ApoE-LNPs, mice were anesthetized with a mixture of xylazine/ketamine (i.p.) and perfused with cold phosphate-buffered saline (PBS) at pH 7.4 and then with 4% paraformaldehyde (PFA) in PBS. Brains were post-fixed in PFA overnight at 4 °C and then put in 30% sucrose for two days, then frozen in isopentane solution and stored at −80 °C. 30 μm-thick brain sections were cut on a cryostat, sections were processed for antigen retrieval (10 min at 85°C in the oven with 100 mM citric acid at pH 6.0), preincubated for 1 h in blocking buffer (5% normal goat serum, 0.1% Triton X-100 in PBS) and incubated overnight at 4 °C with primary antibody (mouse anti-Human Nuclear-Cy3 conjugated, MAB1281, Sigma, 1:100) to label human GBM2 cells. After washing, Hoechst nuclear staining was added (1:300) and the sections were mounted with Vectashield antifade mounting without DAPI (Vector Laboratories). *See “Imaging” methods in Supplemental Information*.

### RNA sequencing and qPCR

RNA isolation and extraction for qPCR were performed with TRIzol (Invitrogen) following manufacturer’s instructions. Reverse transcription and SYBR green-based qPCR for miRNA was performed with miScript II RT kit and primers (Qiagen), human U6 was used as housekeeping gene. <u>Transcriptomics (poly A)</u>. After transfection with the pool or scrambled RNA (n=3 samples per group) GBM cells were seeded at 80000 cells/well in multiwell-24 plates and kept in non-adherent conditions (i.e., in complete medium without Matrigel coating, to allow spheroids formation) for 3 days, then processed for RNA extraction using RNeasy kit (Qiagen). For *in vivo* transcriptomics, GBM xenotransplanted mice were sacrificed 2 weeks (w) after intratumor injection of pool- or scramble-loaded ApoE-LNP (*see “*animal procedures”). Brains were harvested and cut with a slicer matrix kept on ice; 2 mm coronal slices were visually inspected and tumoral tissue was withdrawn with brain punches (Integra LifeSciences) and fresh frozen. Tissue was lysed in RNA extraction buffer (RNeasy kit, Qiagen), homogenized by sonication and processed for RNA extraction (RNAeasy kit, Qiagen). RNA quality was assessed with Bioanalyzer (Agilent). 100 ng of RNA per sample were used to prepare libraries (Illumina Stranded mRNA Prep) and sequenced with a NovaSeq 6000 (Illumina). <u>Small RNA sequencing.</u> RNA was extracted with TRIzol and 1 µg of RNA for each sample was used for library preparation with Nextflex Small RNA-Seq Kit (PerkinElmer, #NOVA-5132-06) with the following modifications: an amplification step on the fragments was carried out with 18 PCR cycles. The quality of the library was assessed by High Sensitivity DNA kit (Agilent, 5067-4626), and bidirectional sequencing was performed with NovaSeq 6000 using a flow cell Reagent Kit v1.5 (Illumina, 20028401). As control, a human postmortem cerebellum and prefrontal cortex small-RNA dataset were used from ^26^, (BioProject PRJNA752352 – accession number: GSE181520). *See “Data Analysis” in Supplemental Information*.

### Proteomics

<u>Samples preparation</u>: GBM1 and 2 were cultured and transfected as above; 24h after transfection cells were detached and seeded at 80000 cells/well in multiwell-24 plates, and kept in non-adherent condition for 6 days. Protein extracts from 5 independent replicates per GBM/per condition, were prepared with ice-cold RIPA buffer containing protease inhibitors (Complete mini EDTA-free, Roche) and sodium orthovanadate (phosphatase inhibitor). *See detailed LC/MS materials and methods in Supplemental Information*.

## Data Availability

Transcriptomics and proteomics datasets generated in this study are available upon request.

## Acknowledgments

The authors wish to thank M. Ghibaudi and F. Macchi for setting up the initial xenotransplantation experiments; M. Morini, E. Petrini and D. Vozzi and the technical staff at IIT’s Animal, Neurofacility and Genomics facility for excellent support. We also wish to thank P. Malatesta at the IRCCS San Martino Hospital in Genoa and all the colleagues at the Neurobiology of miRNA lab of IIT for advice and discussion. We apologize to those colleagues whose work could not be cited due to space limitations.

## Author contributions

Designed research and experiments (SR, PD, DDPT). Performed experiments (SR, SS, RCP, MS; SB, LLR, LG, RP, CB, CS, ALV, MPE, ALP). Data acquisition and analyses (SR, SS, RCP, MS, CB, CS, KT, ALV). Drafted the manuscript (SR, SS, DDPT). Supervision (KT, AA, PD, TF, DDPT). Provided Glioblastoma cultures along with pathological and clinical data from patients (AD, AB, TF). Acquired funding (DDPT). All authors read, revised, and approved the final version of the manuscript.

## Declaration of interests

The pool of eleven miRNA is subject of the Italian priority patent application n. IT102016000093825 by De Pietri Tonelli and Pons-Espinal “A miRNAs Pharmaceutical Composition And Its Therapeutic Uses”, filed on September 19th, 2016 and by the international Patents: EP 17784688.8; US 16/331546; CA 3037254 and JP 2019-515210. All other authors declare no competing interests.

## Funding

This work was funded by IIT and partly by the grant AIRC-IG 2017 # 20106, by the Project “National Center for Gene Therapy and Drug based on RNA Technology” (CN00000041) to DDPT, and by the project MNESYS (PE0000006) – A Multiscale integrated approach to the study of the nervous system in health and disease (DN. 1553 11.10.2022) to TF funded by the Ministry of University and Research (MUR), National Recovery and Resilience Plan (NRRP) #NEXTGENERATIONEU (NGEU).

