## Supplemental information, Supplemental figures and captions for "Synergic microRNAs suppress human glioblastoma progression by modulating clinically relevant targets"

**CONTENT:**


- **Supplemental materials and methods**
- **References for supplemental materials and methods**
- **Supplemental figures S1-S4**
- **Supplemental Tables captions**

#### SUPPLEMENTAL MATERIALS AND METHODS

**LC/MS proteomics**: Protein lysates were kept 30 min in agitation, collected by centrifuged at 13000 x *g* for 30 min at 4°C, then stored at -80°C. 50 µg of proteins (quantified using the colorimetric Bradford dye-binding method, Bio-Rad kit) per sample were reduced with dithiothreitol (100 mM DTT in 50 mM $NH_4HCO_3$) at 56°C for 30 min, alkylated in 100 mM IAA in 50 mM $NH_4HCO_3$ (30 min at room temperature in the dark), then precipitated with cold acetone (-20°C), overnight. Samples were then centrifuged (14000 x *g* for 30 min at 4°C), the pellets were dried under nitrogen stream and resuspended in digestion buffer (50 mM $NH_4HCO_3$). Protein digestion into peptides was obtained by adding 0.5 µg/ml trypsin and letting (37°C overnight). A desalting procedure was conducted with Pierce C18 Spin columns, peptides were then dried and re-dissolved in 3% acetonitrile with 0.1% formic acid. LC/MS analysis: The volume corresponding to 1.65 µg of peptides was injected on a NanoAcquity chromatographic system. The chromatographic method was set as follow: for the trapping phase, an Acquity C18 column (180 µm x 20 mm, from Waters, Milford, MA,USA) was used and the peptides were loaded at 4.0 µl/min (1% acetonitrile + 0.1% FA) for 4 minutes; the peptides were then moved into a PicoFrit C18 column (75 µm x 25 cm, NewObjective Inc.) and eluted at 300 nL/min with a 2 hours gradient of acetonitrile in water (3% to 45%, both eluents were added with 0.1 % FA); the system was washed with 90% of acetonitrile for 10 minutes and then re-equilibrated at 3% of acetonitrile for 18 minutes. The eluted peptides were analyzed by a TripleTof 5600+ mass spectrometer equipped with a NanoSpray III ion source and operating in data-independent acquisition (DIA) mode, following the SWATH protocol for label free proteomics (Braccia et al., 2019). The instrument operated in positive ion mode with the following settings: ion spray voltage at 2000 V, spray gas 1 at 14, curtain gas at 30, declustering potential at 80 V and source temperature at 90 °C. For protein quantification, the tandem mass (MS/MS) spectra were searched against a modified version of the PanHuman ion library (Braccia *et al.*, 2019), using only no-shared peptides. The following parameters were used: minimum peptide confidence 90%, 50 ppm maximum mass tolerance, 30 minutes maximum RT tolerance, 6 MRM transitions per peptide and modified peptides were not allowed. Data analysis: Quantifications of LC/MS Raw data were normalized using the Most Likely Ratio (MLR) method (Huang et al., 2015) before the statistical analysis. To detect significant changes at protein level, an un-paired, 2 tailed t-test was performed and a total of 351 proteins for GBM1 and 462 proteins for GBM2 were found to be significantly (p value < 0,05) altered.

**Lipid Nanoparticles (LNPs).** Microfluidics (Nanoassemblr, Precision NanoSystems) was used to mix one volume of 1,2-dioleoyl-3-dimethylammonium-propane (DODAP), 1,2-dipalmitoyl-sn-glycero-3-phosphocholine (DPPC), 1,2-distearoyl-sn-glycero-3-phosphoethanolamine-N-[methoxy(polyethylene glycol)-2000] (DSPE-PEG2k) (Avanti Polar Lipids Inc) and Cholesterol (Merck) in ethanol (DODAP, DPPC, cholesterol, DSPE-PEG2k at 50/10/38.5/1.5 mol ratio) with 4 volumes of citrate buffer (pH 3.9, 50 mM) containing the miRNAs or scramble RNA (Dharmacon), or a FAM-labeled siRNA (Merck) at the final lipid/RNA ratio of 39:1 (w/w), for the *in vitro* formulation, or 20:1 (w/w) for the *in vivo* one. Total flow rate was 2 ml/min. LNPs were dialyzed for 16-20 h against 500x volume of PBS pH 7.4, then sterilized through 0.22 µm PVDF syringe filters (Millipore). For *in vivo* experiments LNPs were concentrated by centrifugal ultrafiltration units (Amicon Ultra 4, 30kDa MWCO, Millipore). LNP coating with ApoE3: 1 mg of human ApoE3 peptide (Peprotech), every 140 mg lipids, was incubated for 5 min. at 37°C. Diameter, polydispersity index (PdI) and z-potential of LNPs were measured using a Nano-ZS Zetasizer (Malvern) in deionized water. RNA encapsulation efficiency was determined by Ribogreen assay (Thermofisher) following the manufacturer protocol, with a plate reader (Tecan) before and after the lysis of LNPs with Triton X (0.5% w/w). Cryo-electron microscopy of LNPs was done with FEI Tecnai G2-F20. LNPs *in vitro* transfection efficiency: U87MG cells (5x104 cells/well) were seeded into 8 chambered cover glass system for confocal imaging (Lab Tek II, Thermo Scientific) or $8x10^4$/well in multiwell-24 for flow-cytometry analysis and kept in complete medium for 24h. Cells were then treated with LNP loaded with either FAM-labelled siRNA (final siRNA concentration100nM) for 2 or 5 hours and then either fixed with 4% PFA and stained with WGA and DAPI for imaging or analyzed with a BD-FACS aria III cytofluorometer. For qPCR, PD-GBM ($8x10^4$/well) were seeded in multiwell-24 and kept in complete medium for 24h, then treated for

5h with LNP loaded with either miRNA mimics or scrambled RNA (final concentration was 25 nM of each miRNA mimic and 250 nM of scrambled RNA).

**Imaging.** Live cell Imaging**:** Cultured cells were imaged with Incucyte® (Sartorius) and analyzed with Scratch Wound software module, using relative wound density (%) and wound confluence (%) parameters to quantify invasion and migration, respectively, as recommended from vendor. Bright field imaging: 3D spheroids in hydrogel were imaged with a 5X objective in a Leica DMI6000 inverted microscope. Spheroid core area was quantified with Fiji ImageJ software and the relative radius is calculated; the external perimeter formed by spheroids protrusions was selected and the spheroid core area subtracted to obtain the invasion area of each spheroid. Fluorescence imaging (*in vivo* biodistribution): A 10X objective of a microscope Olympus BX51 equipped with the Neurolucida stage and software (MBF Bioscience) was used to acquire an overall visualization of tumor cells (stained with HuNu) in the striatum, and the distribution of FAM-labeled siRNAs (green signal) inside the tumor mass. Then, five images were captured around the injection site of FAM-labeled siRNAs with Nikon A1 confocal microscope at 40x objective and finally some images were taken with 60x objective to observe the localization of FAM-labeled siRNAs inside the tumor cells at higher resolution. IVIS imaging: To evaluate tumor growth *in vivo*, D-Luciferin potassium salt (150mg/kg, PerkinElmer) was intraperitoneally injected into all mice, 10 min after animals were anesthetized and in vivo bioluminescence imaging (BLI) of the tumor growth was captured using the IVIS Spectrum Imaging System (PerkinElmer). Regions of interest (ROI) were defined using Living Image software and total photons flux (p/s) quantified once a week.

**Data Analysis.** Small RNA sequencing: Illumina raw data and the small-RNA FastQ files were processed for adapter content using Cutadapt (version 4.9); untrimmed reads whose coverage did not capture adapter content were merged with the Cutadapt output. The composite files were processed for tetramers, as per library manufacturer instructions, and quantified with miRge3.0, which incorporates the miRNA Transcriptomic Open Project (miRTOP) mirGFF3 standardization of isomiR-miRNA reporting. Outputs were normalized to obtain TPM expression relative to each sample, for each detected miRNA species. TPM values for the 11 miRNAs were selected and plotted into a heatmap. Poly(A) mRNA sequencing, raw data were processed with CLC Genomic Workbench (Qiagen) to perform differential expression analysis (pool vs scramble) and DEG with p value <0.05 were selected. For *in vivo* transcriptomics data, given the presence of mouse tissue contamination in the tumor samples, a human+mouse hybrid reference genome was created (both species' genomes concatenated) and used for alignment; then, reads mapping on more than one species were filtered out and the remaining 42155 human genes used for DEG analysis. Gene ontology (GO) enrichment: Significantly down-regulated protein lists were analyzed separately on DAVID Database V6.8 (https://david.ncifcrf.gov); the whole human genome was used as background. DAVID algorithm uses EASE Score Probability (a modified Fisher Exact P-value) for calculating a P value for gene enrichment; The GO terms were listed from the most significant using Benjamini-corrected P values. PPI analysis and k-means clustering was performed with the STRING software (https://string-db.org) (Szklarczyk et al., 2023). TCGA database: GlioVis Data Visualization Tool for Brain Tumor Datasets (http://gliovis.bioinfo.cnio.es) was employed to analyze and download mRNA expression data encoding the 8 downregulated genes. TCGA RNAseq dataset TCGA_GBMLGG, was used for the correlation analysis of the genes of interest with glioma grade and patient survival. Cell type Cluster analysis: Cluster markers and cell type interpretations data from (Bhaduri et al., 2020) were employed to obtain the lists of marker genes for each cluster. *Proteomics*: Non-significant marker genes were filtered out selecting only adjusted p-values lower than 0.05; then protein-coding gene lists obtained by proteomics data for both PD-GBM (i.e.: total proteome list in ctrl-treated samples; significantly downregulated protein list; significantly up-regulated protein list) were compared with each cluster's marker genes list to obtain the number of genes in common. Non-relevant cell type clusters (B Cells, Dividing B Cells, Red blood cells) were removed from the analysis. The analysis was performed with R-Statistics. *In vitro Transcriptomics:* normalized counts were obtained with DESeq2, and cell clusters markers were used to create custom gene sets to be employed for gene set enrichment analysis with GSEA software. MiRNA-target prediction analysis was performed with DIANA tools (DIANA-microT 2023 webserver (Kavakiotis et al., 2022); validated miRNA-target interactions were retrieved from miRTarBase (Huang et al., 2022).

### SUPPLEMENTAL FIGURES S1-S5

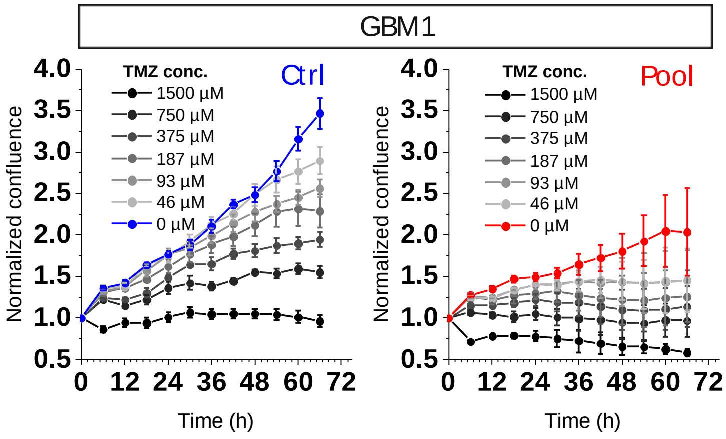


**Supplemental Figure S1)** GBM Growth in presence of TMZ (as determined by cell confluence with automated time-lapse imaging): PD-GBM1 transfected with miRNA pool or scrambled RNA (ctrl) and treated with different TMZ concentrations (1:2 dilutions ranging 1500 to 46 µM) for 72 h. Data expressed as confluence normalized on time 0 (24 h after transfection, i.e. starting of TMZ treatment); mean ± St.Err. of n=1 experiment in triplicates.

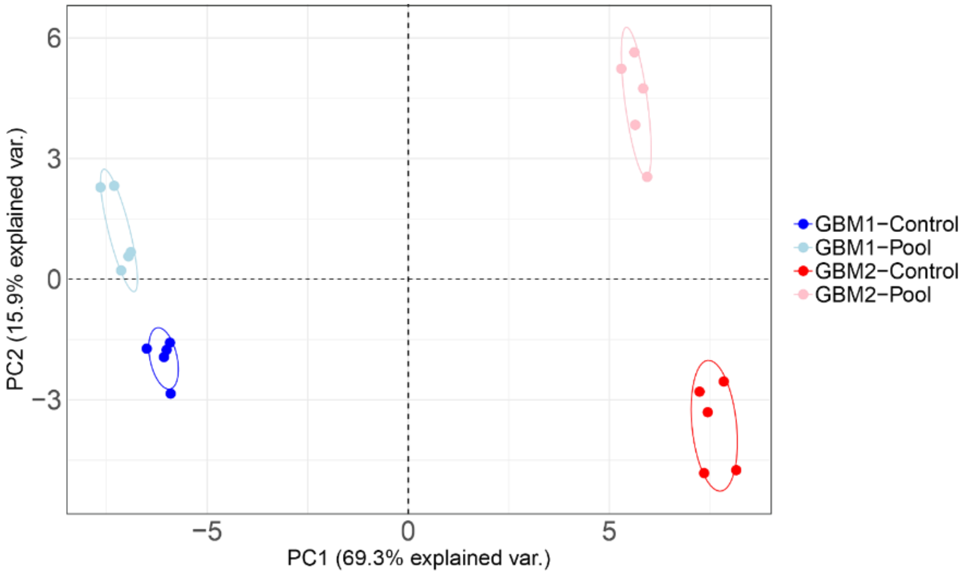


**Supplemental Figure S2)** Principal Component Analysis (PCA) of the significantly altered proteome in scrambled- or pool- transfected GBM1 and GBM2 (n=5 samples/condition per each PD-GBM, 6 days after transfection). Analysis was performed in R. Variables used were the proteins found significantly altered by t-test between treated or non-treated samples. Missing values were removed to process only proteins found in all samples.

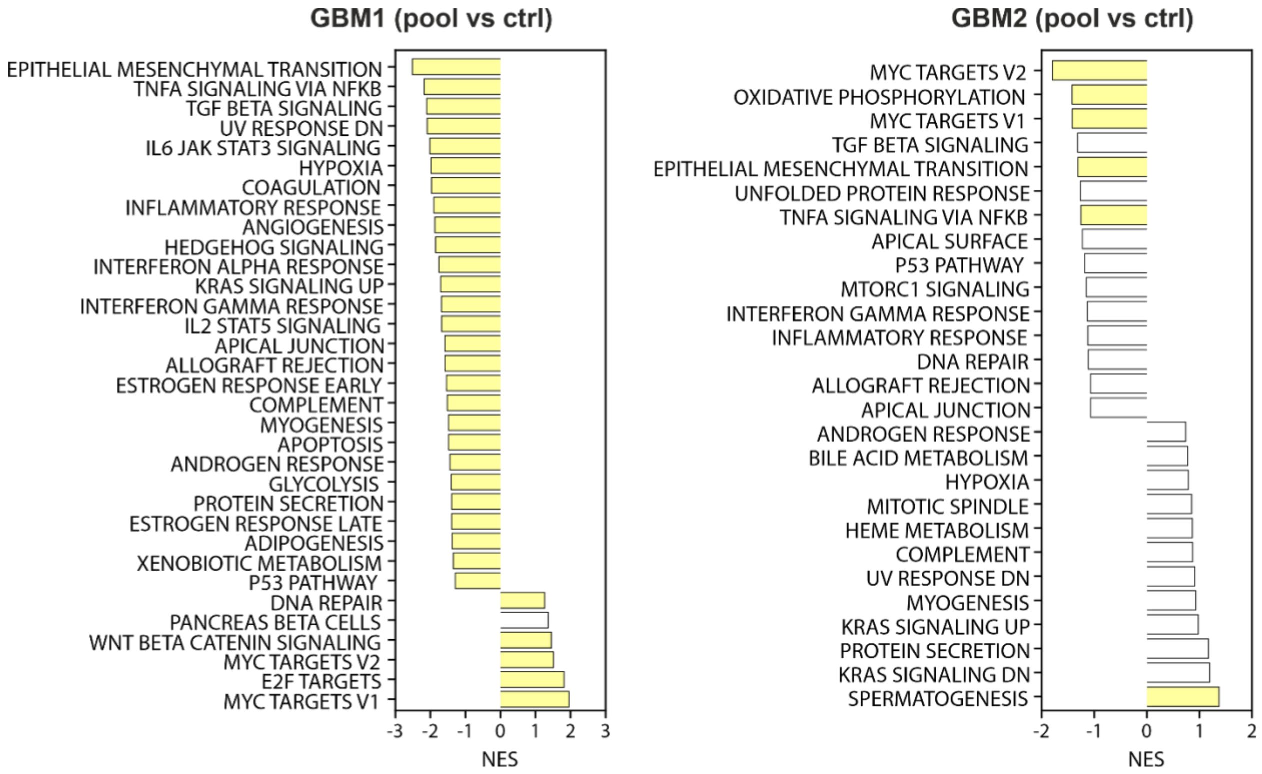


**Supplemental Figure S3)** GSEA pathway analysis of pool vs ctrl-treated GBM1 and GBM2. Negative NES (Normalized Enrichment Score) indicates enrichment in ctrl (scrambled-transfected) samples (i.e. they are down-regulated by the pool), and vice versa. Statistically significantly pathways (NOM q-value <0.05) are filled in yellow.

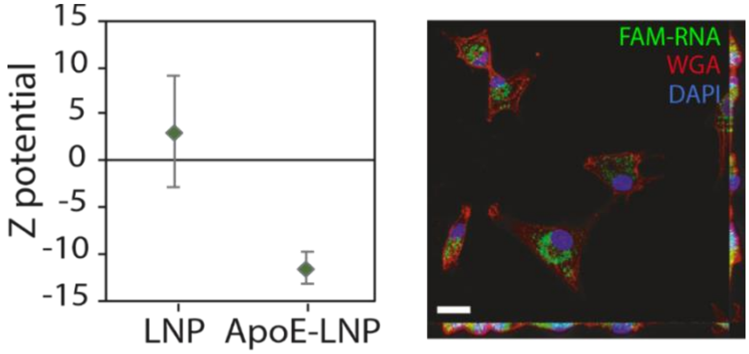


**Supplemental Figure S4) A)** Z potential measurement of LNP with or without ApoE coating (mean ± St.Dev) (n=4 batches measurements). **B**) Representative confocal image of U87 cells, 5h after treatment with 100nM FAM-siRNA loaded in ApoE-LNP. Note the lack of co-localization between RNA (FAM, *green*) signal, the cell surface marker WGA (Alexa 633, *red*) and nucleus (DAPI, *blue*). Maximum projections of the (12 µM-thick) z-stacks are shown on the right and bottom of the image to visualize that FAM-siRNA is inside the cells (scalebar 20 µm).

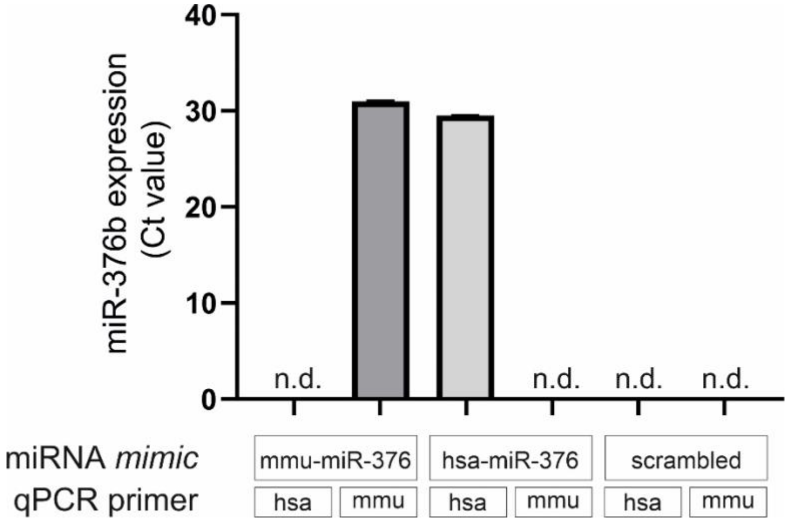


**Supplemental Figure S5)** QPCR analysis of GBM2Luc+ cells transfected with either mmu-miR-376b (mouse-mimic), hsa-miR-376b (human-specific mimic) or scrambled control mimic, and tested for the expression of miR-376b using either human- (hsa) or mouse- (mmu) specific primers. Data are expressed as ct values ±St.Dev (n=3 technical replicates); ct values higher than 35 were plotted as not detected (n.d.). Note that both mouse and human primers are able to detect the overexpression of exogenous mimics in specie-specific way. In scrambled-treated cells, endogenous miR-376b was undetectable using these qPCR settings (in line with our small RNAseq data, Fig. 1B).

**SUPPLEMENTAL TABLES – captions**

**Supplemental Table 1A**: TPM values from small RNA seq of patient-derived glioma cells (GG3, grade III; GBM1, GBM2 grade IV) and healthy brain samples (Hipp: hippocampus; PFC: prefrontal cortex)

**Supplemental Table 1B**. Statistics of miRNA expression from small RNA seq of: patient-derived glioma cells (GG3, grade III; GBM1, GBM2 grade IV) versus healthy brain samples (Hipp: hippocampus; PFC: prefrontal cortex). All significant values are highlighted in yellow (p-value<0.05, t-test)

**Supplemental Table 2A.** Proteomics results: significantly altered proteins in GBM1 upon miRNA pool transfection (pool vs scramble, n=5 biol. replicates per group)

**Supplemental Table 2B**. Proteomics results: significantly altered proteins in GBM2 upon miRNA pool transfection (pool vs scramble, n=5 biol. replicates per group)

**Supplemental Table 2C.** GO enrichment analysis results (DAVID) of down-regulated proteins in GBM1 upon miRNA pool transfection. (In bold significantly enriched pathways, Benjamini-adjusted p-value <0.05).

**Supplemental Table 2D.** GO enrichment analysis results (DAVID) of down-regulated proteins in GBM2 upon miRNA pool transfection. (In bold significantly enriched pathways, Benjamini-adjusted p-value <0.05).

**Supplemental Table 3A.** Transcriptomics results: significantly altered proteins in GBM1 upon miRNA pool transfection (pool vs scramble, n=3 biol. replicates per group)

**Supplemental Table 3B**. Transcriptomics results: significantly altered proteins in GBM2 upon miRNA pool transfection (pool vs scramble, n=3 biol. replicates per group)

**Supplemental Table 3C.** GO enrichment analysis results (DAVID) of down-regulated transcripts in GBM1 upon miRNA pool transfection.

**Supplemental Table 3D.** GO enrichment analysis results (DAVID) of down-regulated transcripts in GBM2 upon miRNA pool transfection.

**Supplemental Table 4.** Correlation between expression of the 8 pool targets and glioma grade from TCGA LGG GBM dataset, including grade II, III and IV gliomas. Statistics generated with Gliovis tool https://gliovis.bioinfo.cnio.es/.
